# ToxCastLite: A portable semantic evidence graph linking in vitro bioactivity, in vivo toxicity, and exposure-use context

**DOI:** 10.64898/2026.05.16.724895

**Authors:** Arif Dönmez, Oleksiy Nosov, Kristina Heck, Axel Mosig, Ellen Fritsche, Katharina Koch

## Abstract

**Motivation:** The ToxCast database is a valuable resource for computational toxicology and new approach methodologies (NAMs), but the approximately 100 GB MySQL distribution is difficult to use for portable local analysis and cross-domain evidence mining. Many practical questions concern chemicals, in vitro bioactivity, in vivo toxicological evidence, and exposure-relevant product-use context rather than raw database keys.

**Results:** We present ToxCastLite, a portable semantic evidence-access system that combines assay-scoped SQLite databases with a compact RDF layer for GraphDB-based querying. The system streams large ToxCast/invitrodb MySQL dumps into curated SQLite profiles, reducing the footprint to approximately 3 GB for focused use cases such as developmental neurotoxicity. Dense numerical evidence, including concentration–response rows, remains in SQLite, while the RDF projection exposes linked semantic entities such as chemicals, assays, endpoints, model results, potency parameters (*AC*_50_), and MC6 quality flags. We further extend the graph with CPDat v4.0 product-use and functional-use evidence and ToxRefDB v3.0 in vivo toxicity evidence, including processed studies, point-of-departure records, effect summaries, and observation summaries. These layers are linked through DSSTox Substance Identifiers, enabling integrated queries across NAM bioactivity, curated animal-study evidence, and exposure/use context. A Streamlit prototype supports exploration through a locally deployed LLM that translates natural-language questions into SPARQL, grounded by a versioned RDF schema to reduce hallucination risk. Case studies in developmental neurotoxicity demonstrate how ToxCastLite identifies concordance between high-confidence in vitro DNT activity and positive in vivo apical evidence, detects in vitro DNT activity beyond available DNT-specific in vivo evidence, and prioritizes chemicals where NAM signals, ToxRefDB evidence, and CPDat product-use context intersect. For selected results, users can drill down from the semantic graph to the underlying SQLite records and retrieve concentration–response curves for expert inspection without manually writing SQL or SPARQL.

**Availability:** Project website at toxcast-lite.github.io/.

## Introduction

The ToxCast database [1; 2; 3] contains high-throughput in vitro screening data, assay metadata, and concentration– response model results, but its distribution as a large relational MySQL resource limits portable and lightweight local analysis. In practice, many users require focused evidence retrieval around chemicals, endpoints, bioactivity profiles, quality flags, in vivo evidence, and exposure-relevant product-use contexts rather than direct interaction with highly normalized relational structures.

ToxCastLite addresses this challenge through a layered architecture that converts periodic ToxCast/invitrodb MySQL dumps into assay-scoped SQLite evidence stores combined with a semantic RDF layer. The design intentionally separates dense numerical evidence from semantic discovery structures. SQLite stores large concentration–response tables and numerical outputs efficiently, while RDF and GraphDB expose discoverable semantic objects and retrieval handles for querying [4]. This makes it possible to query chemicals, assay lists, endpoints, model results, potency parameters such as *AC*_50_, and MC6 quality flags without flattening or discarding the underlying source structure.

In the extended implementation presented here, ToxCastLite also integrates ToxRefDB v3.0 as an in vivo apical evidence layer and CPDat v4.0 as a product-use and functional-use context layer [5; 6]. ToxRefDB contributes processed animal-study records, point-of-departure evidence, effect summaries, and observation summaries, while CPDat contributes functional-use, list-presence, and product-use summaries. These evidence layers are linked through DSSTox Substance Identifiers, enabling cross-domain queries that connect NAM-derived in vitro bioactivity with curated in vivo toxicity evidence and exposure-relevant use contexts.

The framework is designed with transparency, auditability, and reproducibility in mind, making it particularly relevant for regulatory toxicology settings where expert review and traceability are essential. This traceability is facilitated by maintaining a transparent evidence chain: high-level semantic queries can be linked back to the corresponding SQLite records, concentration–response tables, model outputs, quality flags, ToxRefDB evidence objects, and CPDat use-context summaries. Query histories and generated SPARQL can further provide a verifiable audit trail of the discovery process.

## Implementation and features

ToxCastLite combines portable relational evidence stores with a compact semantic access layer. The system is organized around five linked components: assay-scoped SQLite extraction from ToxCast/invitrodb, RDF projection of in vitro bioactivity evidence, CPDat product-use and functional-use context integration, ToxRefDB in vivo evidence integration, and a local query interface with drill-down to dense numerical and source-level records.

### Assay-scoped SQLite evidence stores

The SQLite builder is implemented in Python and processes the large ToxCast database (invitrodb) [3] MySQL dumps in a streaming fashion to avoid loading the full dump into memory. Instead of importing the complete source distribution, the builder retains selected table profiles and prunes the resulting database to curated assay-list categories (Table 1). These scopes cover endocrine, developmental, neurotoxicity, immunotoxicity, cardiotoxicity, mitochondrial, cytotoxicity, and non-mammalian vertebrate assay domains.

**Table 1.** Currently available assay-list scopes for assay-scoped ToxCastLite builds.

| ID | Assay-list scope | Description |
| --- | --- | --- |
| 79 | Cytotoxicity Burst | Assays used to define the cytotoxicity burst region |
| 151 | ToxCast ER Pathway Model | Estrogen receptor pathway model assays |
| 152 | ToxCast AR Pathway Model | Androgen receptor pathway model assays |
| 153 | Thyroid Bioactivity | Assays related to thyroid bioactivity |
| 154 | Steroidogenesis Bioactivity | Assays related to steroidogenesis |
| 155 | Developmental Toxicity | Assays associated with developmental toxicity |
| 156 | DNT-IVB | Developmental neurotoxicity in vitro battery assays |
| 157 | Immunosuppression | Assays associated with markers of immunosuppression |
| 158 | Cardiotoxicity | Assays associated with cardiotoxicity |
| 159 | Non-mammalian Vertebrate | Assays associated with non-mammalian vertebrate species |
| 160 | Mitochondrial Toxicity | Assays associated with mitochondrial toxicity |

The build process consists of four steps: selected MySQL `CREATE TABLE` statements are translated into SQLite-compatible definitions; selected `INSERT` statements are parsed and imported in batches; assay-list pruning removes rows outside the requested endpoint scope while preserving the join paths needed to connect chemicals, samples, assays, endpoints, model results, and quality flags; and the database is compacted using `VACUUM`. The final compaction step is important because pruning removes rows logically, whereas SQLite releases unused pages physically only after vacuuming.

Dense numerical tables, including raw and normalized concentration–response records, remain in SQLite. This keeps the evidence store directly usable from Python, R, command-line SQLite, or the semantic application layer while avoiding unnecessary duplication of measurement-level data.

### RDF projection of ToxCast bioactivity evidence

The RDF projection is generated from the assay-scoped SQLite database and emitted as Turtle. Rather than duplicating the relational schema one-to-one, it exposes the retained evidence store as typed graph entities organized around chemical identity, assay hierarchy, endpoint membership, concentration–response evidence handles, model results, potency parameters, and downstream quality flags.

Typed relations preserve the original invitrodb evidence chain, linking chemicals to samples, samples to assay results, endpoints to assay-list scopes, and model/flag objects to their corresponding endpoints and retrievable concentration–response tables. Detailed class coverage and graph connectivity are summarized in Table 2. In this design, GraphDB acts as a semantic index over the evidence store [4], whereas SQLite remains the authoritative storage layer for dense numerical data.

**Table 2.**
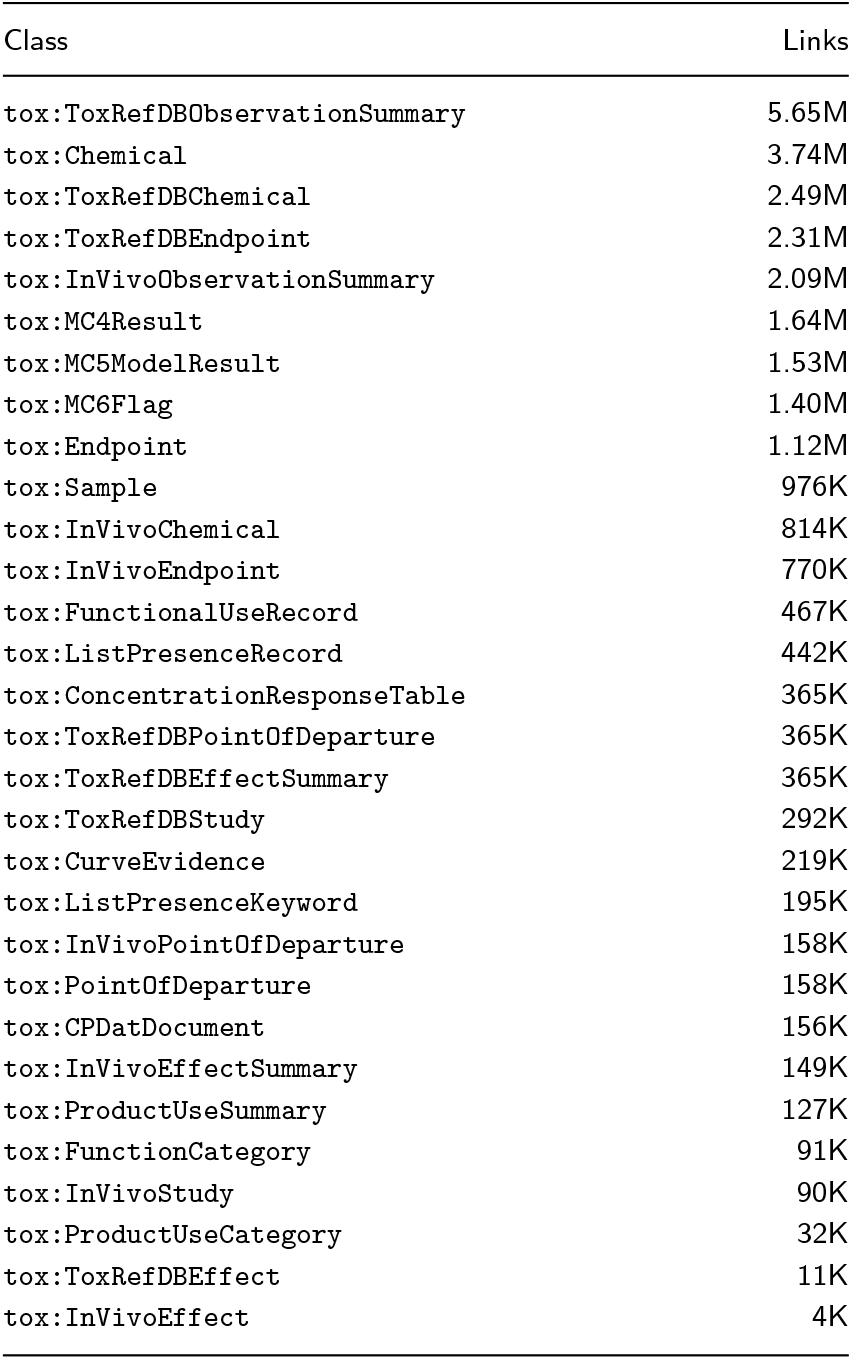
GraphDB class-relationship connectivity for the integrated DNT-scoped evidence graph with CPDat and ToxRefDB extensions. Values are rounded as displayed in the GraphDB class-relationships view and describe link connectivity rather than simple relational row or instance counts. The table shows the 30 most connected classes.

A central design choice is that individual concentration– response points are not materialized as RDF triples. Instead, the graph contains `tox:ConcentrationResponseTable` entities. Each entity represents a retrievable concentration–response table for a chemical/sample–endpoint combination and stores SQLite identifiers such as endpoint ID and sample ID. Model-result entities such as `tox:MC5ModelResult` expose hit-call and potency-related fields, including *AC*_50_ where available, while flag entities such as `tox:MC6Flag` preserve quality and decision-support information. This enables graph-level filtering followed by local numerical drill-down.

### CPDat product-use context integration

CPDat v4.0 [8] is integrated as a complementary product-use and functional-use context layer. CPDat distinguishes three document-derived evidence streams: product-composition records, functional-use records, and list-presence records. Composition documents provide information on chemicals reported in products, functional-use documents describe roles that chemicals may serve in products or processes, and list-presence documents capture broader evidence that a chemical occurs on curated use-related lists. These evidence streams are organized using controlled vocabularies for product-use categories, functional categories, and list-presence keywords.

The CPDat workflow imports the v4.0 CSV and metadata files into SQLite and projects selected product-use evidence streams into RDF. The projection preserves CPDat’s distinction between product-composition, functional-use, and list-presence evidence, linking contextual records to ToxCast chemicals through DSSTox Substance Identifiers where available. Full CPDat records remain in SQLite for drill-down and provenance inspection, while the RDF layer supports semantic filtering across bioactivity and use-context evidence. The corresponding CPDat evidence streams and RDF representations are summarized in Table 3.

**Table 3.**
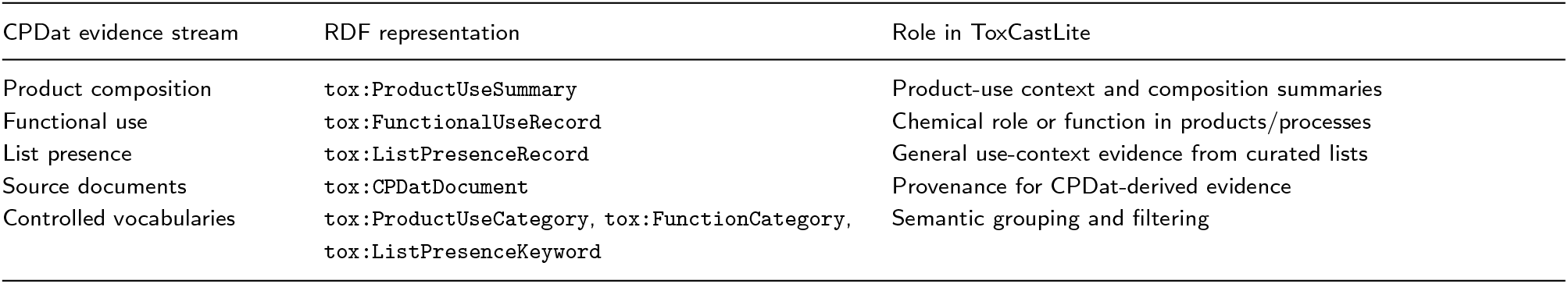
CPDat evidence streams represented in the ToxCastLite semantic layer.

CPDat records are represented as contextual evidence rather than exposure estimates. This preserves a clear distinction between bioactivity evidence, chemical-use evidence, and quantitative exposure modeling. The graph can therefore support combined prioritization queries, such as identifying chemicals active in a selected ToxCast assay domain that also have CPDat functional-use or product-use evidence, while retaining access to the underlying CPDat document provenance.

### ToxRefDB in vivo evidence integration

ToxRefDB v3.0 is integrated as an in vivo apical evidence layer [5; 6]. The public PostgreSQL release was first loaded into a local PostgreSQL instance and then mirrored into a schema-preserving SQLite database. The mirror retains the core ToxRefDB relational structure, including chemicals, studies, treatment groups, doses, endpoints, effects, observation records, guideline profiles, negative endpoint/effect records, and point-of-departure records. This preserves detailed in vivo evidence for local inspection while allowing the RDF graph to expose query-relevant summary entities.

The ToxRefDB RDF projection represents chemicals, processed studies, point-of-departure records, endpoint and effect vocabulary, effect summaries, and observation summaries. ToxRefDB chemicals are linked to ToxCast chemicals through shared DSSTox Substance Identifiers. This creates direct graph paths from in vitro bioactivity and assay-quality evidence to curated in vivo animal-study evidence. For example, a ToxCast chemical can be linked to ToxRefDB studies, LOAEL point-of-departure records, critical-effect summaries, and endpoint observation summaries.

As with the ToxCast and CPDat layers, the projection avoids materializing all dense source records as RDF triples. Highly granular ToxRefDB tables, including detailed dose-treatment group effects, negative effects, and full observation records, remain in SQLite. The RDF layer instead stores semantic summaries and retrieval handles that support discovery, filtering, ranking, and cross-database linking. This design allows users to ask whether an in vitro NAM signal is supported by positive in vivo apical evidence, whether DNT-specific in vivo evidence is absent despite strong in vitro activity, or whether a chemical also appears in product-use contexts relevant to prioritization.

### Local query interface and drill-down

The frontend prototype is implemented in Streamlit and connects to GraphDB and the local SQLite evidence stores. Expert users can submit SPARQL directly. In addition, the interface supports schema-grounded natural-language query generation through a locally deployed model. The model acts as a backend translator rather than a knowledge source: it generates candidate SPARQL from a versioned RDF/SPARQL context that defines the allowed classes, predicates, and query patterns.

This architecture keeps the system deployable in local or institutional environments without external API dependencies. GraphDB is used for semantic discovery, ranking, filtering, and evidence linking. SQLite is used for dense numerical and source-level retrieval. When a SPARQL query returns a `tox:ConcentrationResponseTable` handle, the application retrieves the corresponding concentration–response rows from SQLite and forwards them to an optional R service for concentration–response analysis, plotting, and expert review [7]. Analogously, ToxRefDB and CPDat graph entities retain identifiers that allow users to drill down into the corresponding SQLite records for study, effect, observation, point-of-departure, product-use, or provenance details.

This design supports three user levels. Expert users can submit SPARQL directly; domain users can use controlled query patterns, such as asking for active chemicals within a given assay-list scope or chemicals with positive ToxRefDB evidence; and review-oriented users can drill down from a semantic result to the concentration–response rows, model output, flags, ToxRefDB evidence objects, and CPDat source context that support the result. Thus, the same system can serve as a semantic index, a local evidence browser, and a reproducible bridge between graph-level discovery and numerical, in vivo, and product-use toxicology data.

## Applications

ToxCastLite is intended for users who need to inspect and prioritize toxicological evidence without directly working with the full invitrodb, ToxRefDB, or CPDat relational structures. The primary target users are toxicologists, risk assessors, data stewards, and regulatory scientists in agencies or agency-facing research environments. These users may understand the biological and regulatory questions, but may not be familiar with MySQL or PostgreSQL schemas, RDF vocabularies, SPARQL syntax, or the detailed tcpl and ToxRefDB processing hierarchies.

The system is therefore designed as a semantic access layer rather than as an automated decision system. It supports evidence retrieval, prioritization, and drill-down while keeping the final interpretation under expert control. In this setting, ToxCastLite acts as a local evidence browser: GraphDB identifies relevant chemicals, endpoints, model results, quality flags, ToxRefDB in vivo evidence, and CPDat product-use context, whereas SQLite stores the dense numerical and source-level evidence needed for expert review.

A typical regulatory user may ask a question such as: *“Which high-potency DNT-active chemicals are associated with children’s products?”* The interface translates the request into a schema-grounded SPARQL query that combines assay-list membership, active model-result objects with potency-related fields, and CPDat-derived product-use context. A more integrated question may ask: *“Which high-confidence DNT-active chemicals have positive ToxRefDB apical evidence and exposure-relevant product-use context?”* This query combines in vitro NAM activity, assay-quality metadata, curated in vivo evidence, and product-use context in a single evidence workflow.

The returned result is not a regulatory conclusion, but a ranked evidence table that can be inspected further. For each chemical, the user can drill down to active endpoints, model fields, quality flags, CPDat context, ToxRefDB studies, point-of-departure records, effect summaries, observation summaries, and the underlying concentration–response records stored in SQLite. This allows the user to evaluate whether a screening-level alert is isolated, supported by positive in vivo apical evidence, or embedded in broader product-use and hazard-evidence contexts.

Additional supported query patterns include identifying chemicals active in a selected assay domain, retrieving active endpoint profiles for a chemical of interest, finding chemicals with complete endpoint coverage within an assay-list scope, linking high-confidence in vitro activity to ToxRefDB point-of-departure or critical-effect evidence, identifying chemicals with strong in vitro activity but no available DNT-specific positive in vivo evidence, and combining ToxCast bioactivity evidence with CPDat functional-use or product-use context. Representative question patterns and the corresponding evidence returned by the system are summarized in Table 4.

**Table 4.**
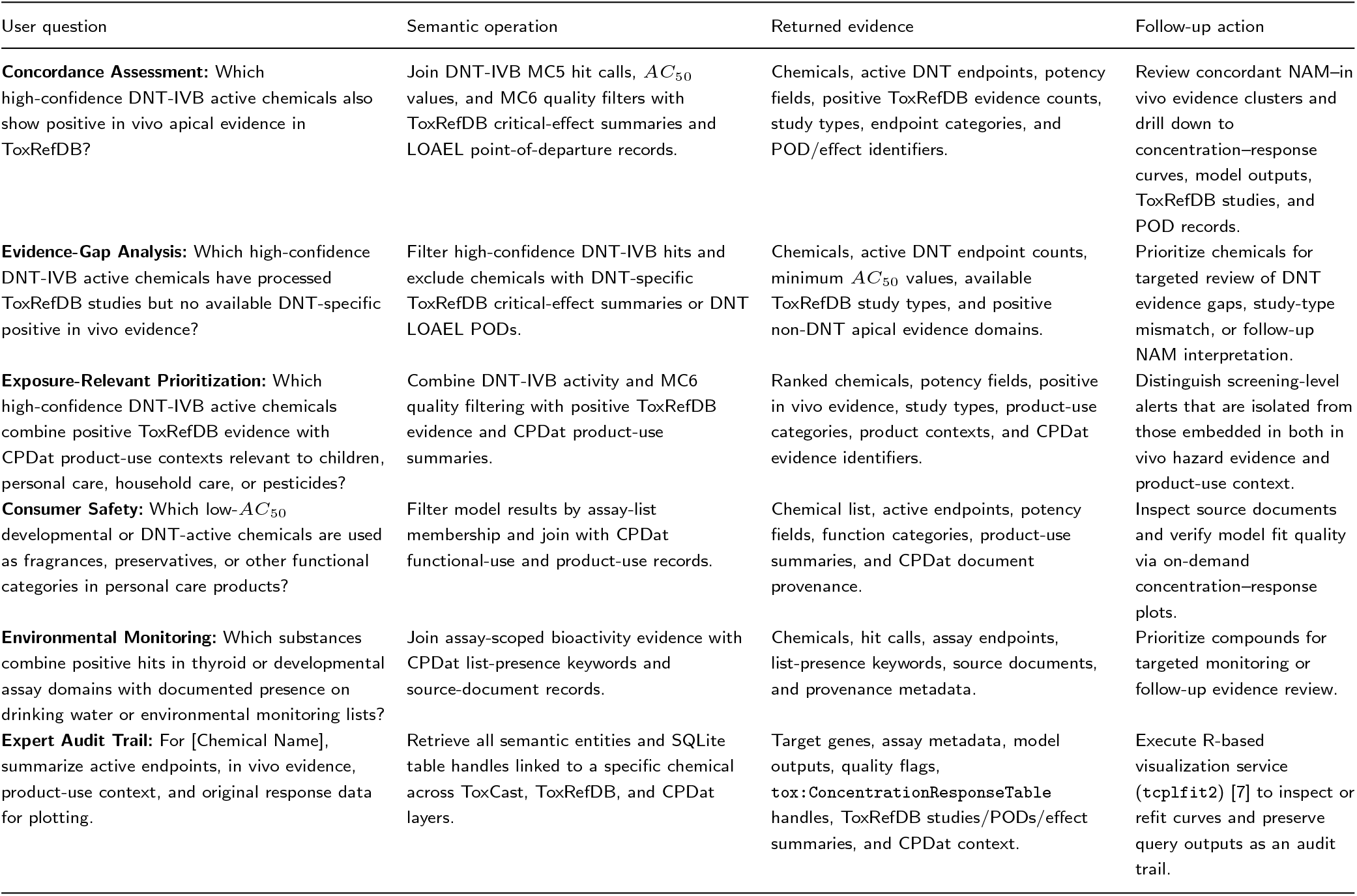
A selection of regulatory and scientific questions supported by ToxCastLite after integration of ToxCast bioactivity, ToxRefDB in vivo evidence, and CPDat product-use context.

Because the system is designed for local deployment, agencies can use ToxCastLite without sending queries or data to external services. The same build scripts, RDF projection, SPARQL context, and SQLite files can be archived with a project, allowing evidence retrieval to be repeated later. Query histories, generated SPARQL, and selected drill-down outputs can be preserved as part of a review record, supporting traceability from high-level semantic results back to the underlying in vitro numerical evidence, curated in vivo evidence, and product-use provenance.

### Case Study 1: Concordance between in vitro DNT activity and in vivo apical evidence

As a first case study, we evaluated whether high-confidence in vitro DNT activity could be linked to positive in vivo apical evidence in ToxRefDB. To mimic the intended user workflow, we entered the following natural-language question into the Streamlit query interface:

> *Which chemicals combine high-confidence in vitro DNT-IVB activity with positive in vivo apical evidence in ToxRefDB?*

In natural-language mode, the interface routed this question to the local schema-grounded LLM translator, which generated a SPARQL query against the integrated ToxCastLite–ToxRefDB graph. The generated SPARQL query was inspected and executed in GraphDB; it is reported in Appendix A.2.1 for reproducibility.

High-confidence in vitro DNT activity was defined by four criteria: a positive MC5 hit call, an AC50 below 10 *µ*M, membership in the DNT-IVB assay list, and the absence of MC6 quality flags. Positive in vivo apical evidence was operationalized conservatively as either a ToxRefDB critical-effect summary with at least one critical effect or a LOAEL point-of-departure record. We restricted the in vivo evidence layer to developmental neurotoxicity (DNT), developmental toxicity (DEV), multigeneration reproductive toxicity (MGR), chronic toxicity (CHR), and subchronic toxicity (SUB) studies.

The translated query identified 136 chemicals that combined high-confidence DNT-IVB activity with positive in vivo apical evidence. These chemicals therefore represent concordant evidence clusters, where NAM-derived bioactivity is not only detectable in vitro, but can also be connected to positive apical evidence from curated animal toxicity studies.

To characterize the biological nature of the concordant in vivo evidence, we further decomposed the associated ToxRefDB effect annotations into broad apical-effect categories. Effect annotations were grouped by their first-level ToxRefDB effect category, and each chemical was counted at most once per category. This avoids overweighting chemicals with many repeated study annotations and emphasizes the breadth of in vivo apical domains represented among the concordant NAM-active chemicals.

Several patterns are apparent from Table 5 and Figure 2. First, multiple chemicals showed very low in vitro AC50 values together with positive in vivo evidence across several ToxRefDB study types. Thiacloprid, Cymoxanil, Pyridaben, emamectin benzoate, and others linked to positive in vivo evidence across developmental, reproductive, chronic, subchronic, and in some cases DNT studies. Second, several chemicals showed broad DNT-IVB activity across many endpoints. For example, abamectin, emamectin benzoate, pyridaben, and fenpyroximate were active across more than 30 DNT-IVB endpoints while also linking to positive in vivo evidence.

**Table 5.**
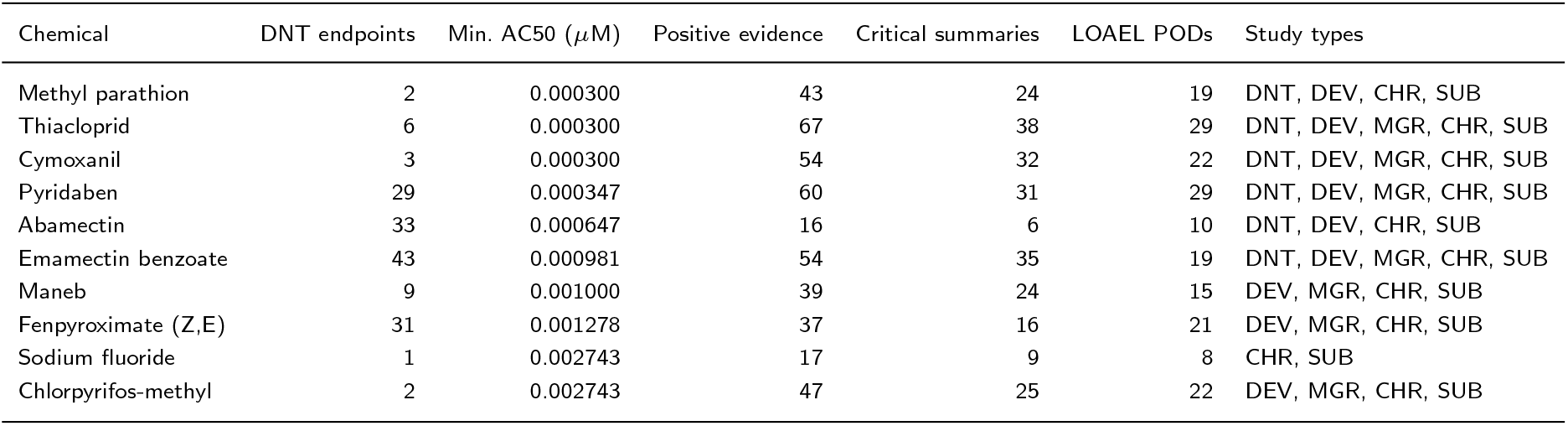
Top high-confidence DNT-IVB active chemicals with positive in vivo apical evidence in ToxRefDB. Positive in vivo evidence was defined as a ToxRefDB critical-effect summary or LOAEL point-of-departure record in DNT, DEV, MGR, CHR, or SUB studies. The “Positive evidence” column counts distinct positive ToxRefDB evidence objects, not independent studies.

| Chemical | DNT endpoints | Min. AC50 (μM) | Positive evidence | Critical summaries | LOAEL PODs | Study types |
| --- | --- | --- | --- | --- | --- | --- |
| Methyl parathion | 2 | 0.000300 | 43 | 24 | 19 | DNT, DEV, CHR, SUB |
| Thiacloprid | 6 | 0.000300 | 67 | 38 | 29 | DNT, DEV, MGR, CHR, SUB |
| Cymoxanil | 3 | 0.000300 | 54 | 32 | 22 | DNT, DEV, MGR, CHR, SUB |
| Pyridaben | 29 | 0.000347 | 60 | 31 | 29 | DNT, DEV, MGR, CHR, SUB |
| Abamectin | 33 | 0.000647 | 16 | 6 | 10 | DNT, DEV, CHR, SUB |
| Emamectin benzoate | 43 | 0.000981 | 54 | 35 | 19 | DNT, DEV, MGR, CHR, SUB |
| Maneb | 9 | 0.001000 | 39 | 24 | 15 | DEV, MGR, CHR, SUB |
| Fenpyroximate (Z,E) | 31 | 0.001278 | 37 | 16 | 21 | DEV, MGR, CHR, SUB |
| Sodium fluoride | 1 | 0.002743 | 17 | 9 | 8 | CHR, SUB |
| Chlorpyrifos-methyl | 2 | 0.002743 | 47 | 25 | 22 | DEV, MGR, CHR, SUB |

**Figure 1.**
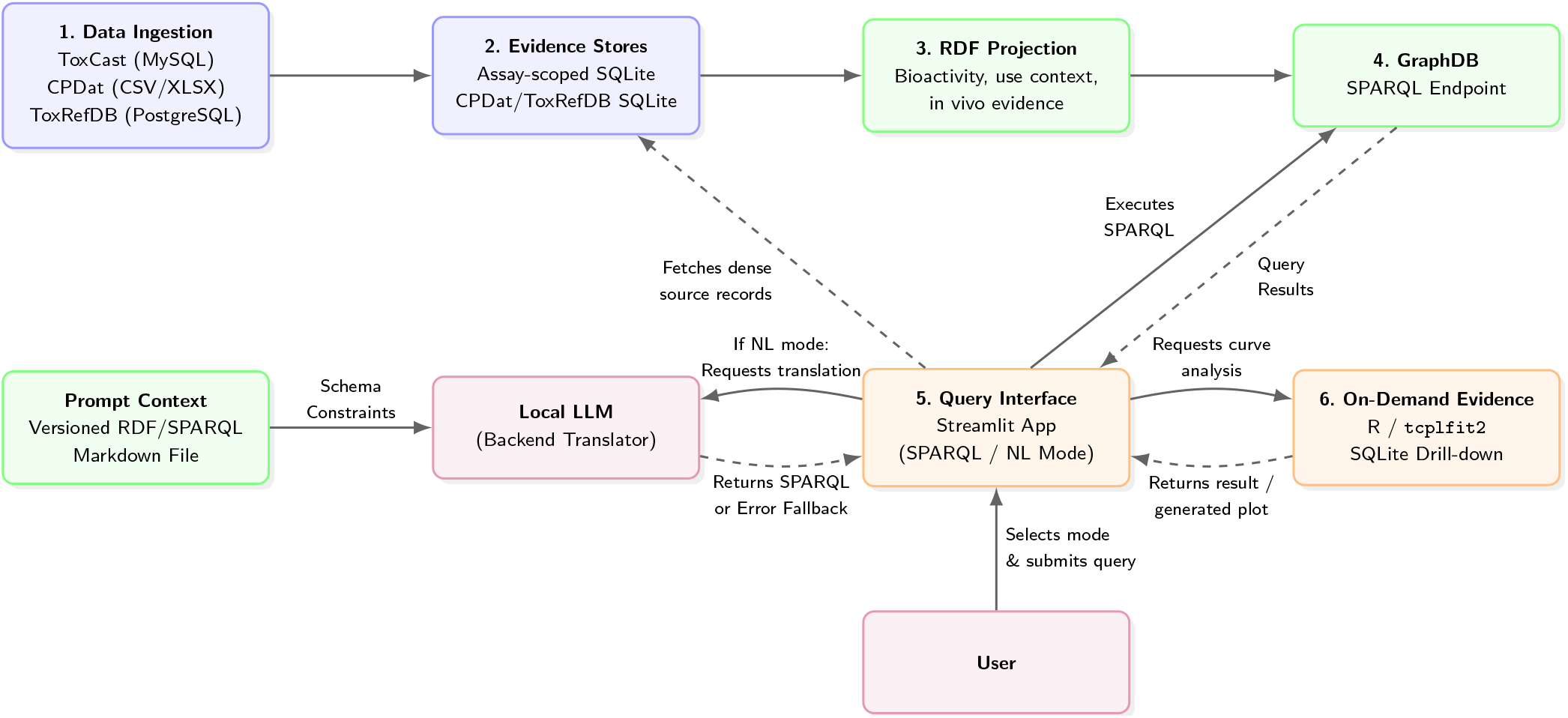
End-to-end workflow of ToxCastLite. Source data from ToxCast/invitrodb, CPDat, and ToxRefDB are converted into portable SQLite evidence stores. The ToxCast layer retains dense concentration–response and model-output data, while CPDat and ToxRefDB retain product-use, functional-use, study, point-of-departure, effect, and observation records for local drill-down. Query-relevant entities are projected into RDF and served through a GraphDB SPARQL endpoint, linking in vitro bioactivity, assay-quality metadata, product-use context, and in vivo apical evidence through shared chemical identifiers. The user interacts with the Streamlit interface in either SPARQL or natural-language mode. In natural-language mode, the application routes the request to a locally hosted LLM acting strictly as a backend translator, constrained by a versioned Markdown RDF/SPARQL context. Successfully generated or directly supplied SPARQL queries are executed against GraphDB, while dense numerical and source-level records are retrieved from SQLite on demand for expert review, curve visualization, and downstream analysis [7].

**Figure 2.**
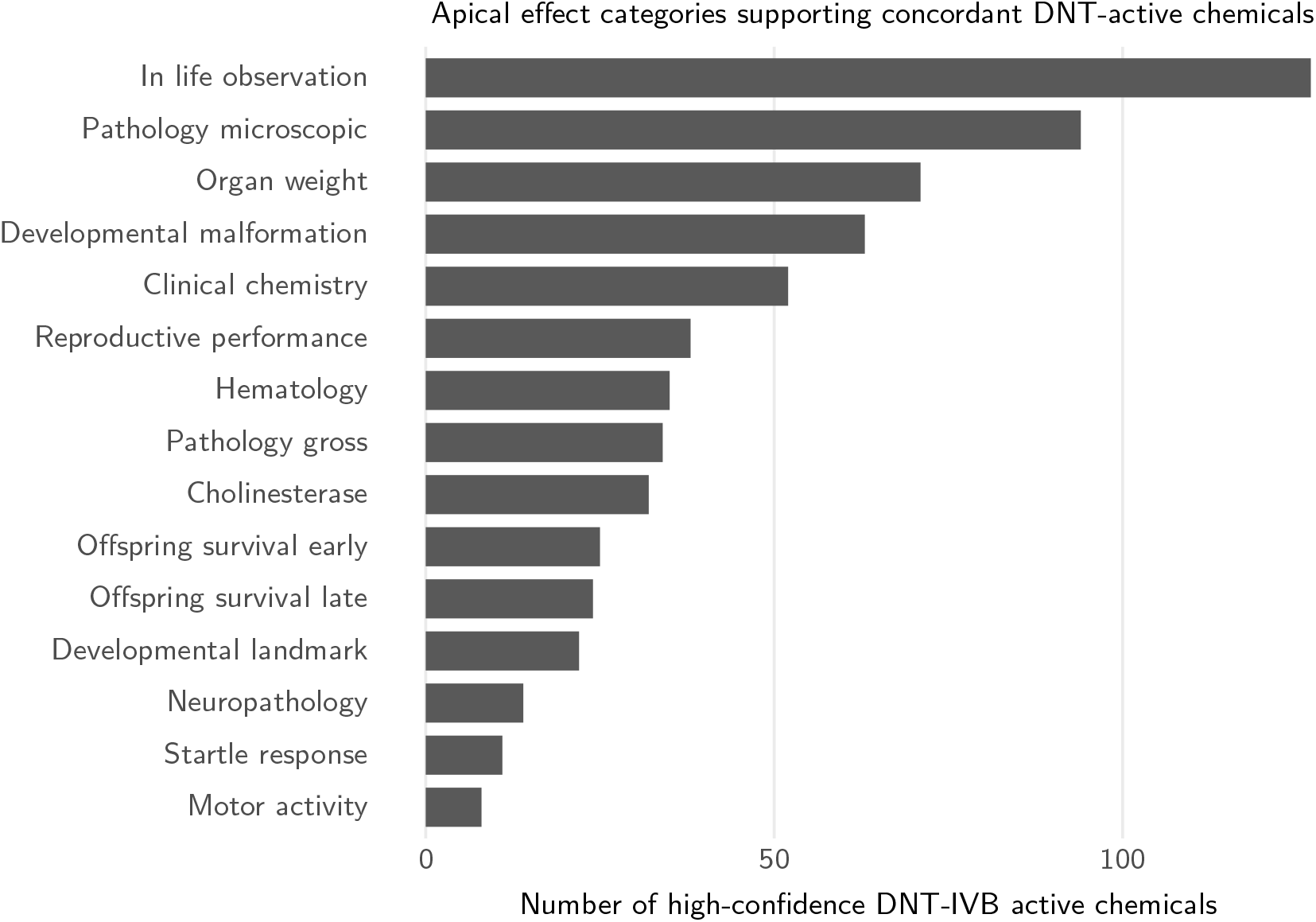
Frequency of ToxRefDB apical effect categories among high-confidence DNT-IVB active chemicals with positive in vivo evidence. Effect annotations were grouped by first-level ToxRefDB effect category, and each chemical was counted at most once per category.

The effect-category distribution shows that the concordant chemicals were not supported by a single narrow in vivo endpoint domain. Instead, positive ToxRefDB evidence covered broad apical categories including in-life observations, microscopic pathology, organ weight changes, developmental malformations, clinical chemistry, reproductive performance, cholinesterase-related effects, and neuropathology. This places the DNT-IVB NAM signals into a broader apical toxicity landscape rather than treating them as isolated screening hits.

This case study illustrates the value of the integrated evidence graph for identifying concordance between NAM-derived in vitro activity and curated in vivo toxicity evidence through a reproducible natural-language-to-SPARQL workflow. Importantly, the analysis does not assume a one-to-one correspondence between individual in vitro endpoints and apical in vivo outcomes. Rather, it identifies chemicals for which independent evidence layers are concordant at the level of prioritization: potent DNT-IVB bioactivity, absence of technical quality flags, and positive apical evidence in ToxRefDB. Such concordant evidence clusters provide a transparent starting point for prioritization, follow-up review, and regulatory interpretation.

### Case Study 2: In vitro DNT activity beyond available in vivo DNT evidence

As a second case study, we evaluated whether high-confidence in vitro DNT activity could identify chemicals for which the available in vivo evidence does not contain DNT-specific positive evidence. To mimic the intended user workflow, we entered the following natural-language question into the Streamlit query interface:

> *Which high-confidence DNT-IVB active chemicals have processed ToxRefDB study evidence, but no available DNT-specific positive in vivo evidence?*

In natural-language mode, the interface routed this question to the local schema-grounded LLM translator, which generated a SPARQL query against the integrated ToxCastLite–ToxRefDB graph. The generated SPARQL query was inspected and executed in GraphDB; it is reported in Appendix A.2.2 for reproducibility.

We used the same high-confidence in vitro definition as in Case Study 1: a positive MC5 hit call, an *AC*_50_ below 10 *µ*M, membership in the DNT-IVB assay list, and the absence of MC6 quality flags. We then required processed ToxRefDB study evidence in DNT, DEV, MGR, CHR, or SUB studies, but excluded chemicals with DNT-specific positive in vivo evidence. DNT-specific positive evidence was defined conservatively as either a DNT critical-effect summary with at least one critical effect or a DNT LOAEL point-of-departure record. For the remaining chemicals, we summarized positive non-DNT in vivo evidence from DEV, MGR, CHR, and SUB studies.

The translated query identified 87 chemicals. These chemicals are not evidence-free: they have processed ToxRefDB study evidence and, in many cases, positive apical evidence in non-DNT study types. However, under the operational definition used here, they lack available DNT-specific positive in vivo evidence. This makes them informative for identifying cases where in vitro DNT-IVB activity extends beyond what is directly represented as DNT-positive evidence in the available in vivo ToxRefDB layer.

Several patterns are apparent from Table 6. First, the absence of DNT-specific positive in vivo evidence does not imply absence of in vivo toxicity evidence. Most chemicals in this subset had positive non-DNT evidence in systemic toxicity domains, and many also linked to developmental, reproductive, or cholinesterase-related apical evidence. At the chemical level, positive non-DNT evidence occurred most frequently in systemic endpoints (82 chemicals), followed by developmental endpoints (37 chemicals), reproductive endpoints (22 chemicals), and cholinesterase-related endpoints (13 chemicals). Second, several chemicals showed broad in vitro DNT-IVB activity despite lacking DNT-specific positive in vivo evidence under the definition used here. Fenpyroximate, for example, was active across 31 DNT-IVB endpoints, while rotenone was active across 33 DNT-IVB endpoints in the full result set. These cases are particularly useful for evidence-gap analysis because they combine strong in vitro DNT activity with broader, but not DNT-specific, in vivo apical evidence.

**Table 6.** High-confidence DNT-IVB active chemicals with processed ToxRefDB study evidence but no available DNT-specific positive in vivo evidence. Positive non-DNT evidence was defined as a critical-effect summary or LOAEL point-of-departure record in DEV, MGR, CHR, or SUB studies.

| Chemical | DNT endpoints | Min. $AC_{50}$ (μM) | ToxRefDB studies | Positive non-DNT evidence | Positive non-DNT domains |
| --- | --- | --- | --- | --- | --- |
| Maneb | 9 | 0.001000 | 6 | 39 | systemic, reproductive, developmental |
| Fenpyroximate (Z,E) | 31 | 0.001278 | 8 | 37 | systemic, developmental |
| Sodium fluoride | 1 | 0.002743 | 4 | 17 | systemic |
| Chlorpyrifos-methyl | 2 | 0.002743 | 7 | 47 | cholinesterase, systemic, developmental |
| Vinclozolin | 2 | 0.002786 | 12 | 73 | systemic, reproductive, developmental |
| Sodium chlorite | 5 | 0.003000 | 1 | 7 | developmental, systemic |
| L-Ascorbic acid | 3 | 0.003000 | 4 | 11 | systemic |
| 2-(m-Chlorophenoxy)propionic acid | 2 | 0.003000 | 3 | 11 | developmental, systemic |
| Glyphosate | 1 | 0.003000 | 7 | 35 | systemic, developmental |
| Tris(methylphenyl) phosphate | 2 | 0.003000 | 1 | 4 | systemic, cholinesterase |

This result should not be interpreted as evidence that these chemicals are negative in vivo for DNT. Rather, it indicates that the available processed ToxRefDB evidence does not contain DNT-specific positive evidence under the conservative operational definition used here. The distinction is important: absence of DNT-specific positive evidence may reflect lack of DNT testing, incomplete endpoint coverage, study-type mismatch, curation boundaries, or true discordance between in vitro and in vivo evidence layers.

The case study therefore illustrates a complementary role for NAMs within a reproducible natural-language-to-SPARQL workflow. Instead of merely reproducing the structure of existing animal-study databases, in vitro DNT assays can highlight chemicals whose bioactivity is not directly captured as DNT-positive in the available in vivo evidence layer, while still being embedded in broader apical toxicity profiles. This supports the use of NAM-derived evidence as a guide for targeted follow-up, evidence-gap analysis, and prioritization.

### Case Study 3: Exposure-relevant prioritization across NAM, in vivo, and product-use evidence

As a third case study, we evaluated whether the integrated evidence graph could prioritize chemicals at the intersection of three evidence layers: high-confidence in vitro DNT activity, positive in vivo apical evidence, and product-use context. To mimic the intended user workflow, we entered the following natural-language question into the Streamlit query interface:

> *Which high-confidence DNT-IVB active chemicals combine in vitro potency, positive ToxRefDB in vivo apical evidence, and CPDat product-use contexts relevant to children, toys, personal care, household care, or pesticide exposure?*

In natural-language mode, the interface routed this question to the local schema-grounded LLM translator, which generated a SPARQL query against the integrated ToxCastLite–ToxRefDB– CPDat graph. The generated SPARQL query was inspected and executed in GraphDB; it is reported in Appendix A.2.3 for reproducibility.

We used the same high-confidence in vitro definition as in the previous case studies: a positive MC5 hit call, an *AC*_50_ below 10 *µ*M, membership in the DNT-IVB assay list, and the absence of MC6 quality flags. Positive in vivo apical evidence was again defined as either a ToxRefDB critical-effect summary with at least one critical effect or a LOAEL point-of-departure record in DNT, DEV, MGR, CHR, or SUB studies. We then required at least one CPDat product-use summary in an exposure-relevant product context, including children’s products, toys, personal care, household or cleaning products, or pesticides.

The translated query identified 61 chemicals that satisfied all three criteria. These chemicals represent a highly prioritized subset in which NAM-derived DNT bioactivity, positive in vivo apical evidence, and product-use context co-occurred in a single queryable evidence workflow. Importantly, CPDat product-use context should not be interpreted as measured exposure or risk. Rather, it provides an interpretable use-context layer that can help prioritize chemicals for follow-up evaluation when combined with bioactivity and in vivo evidence.

To summarize the product-use landscape of this prioritized subset, we grouped CPDat product-use summaries by broad product category and counted each chemical at most once per category. The resulting distribution showed that the largest product-use category was pesticides, followed by personal care, cleaning and safety, and cleaning products and household care. Smaller categories included food and drug contexts. This distribution reflects the combined evidence filters used here: the selected chemicals are not merely DNT-IVB active in vitro, but also have positive in vivo apical evidence and product-use annotations in CPDat.

Several patterns are apparent from Table 7 and Figure 3. First, pesticides formed a prominent product-use category among chemicals that also showed high-confidence DNT-IVB activity and positive in vivo evidence. This is consistent with the strong representation of pesticide-related chemicals in both ToxCast and ToxRefDB. Second, the presence of personal care, cleaning, household, food and drug, and related product-use categories demonstrates that the same evidence graph can extend beyond hazard-oriented data integration and into use-context-aware prioritization. For example, sodium fluoride and sodium chlorite linked DNT-IVB activity and positive ToxRefDB evidence to personal care or cleaning-related CPDat contexts, while diazinon combined in vitro DNT activity, positive in vivo apical evidence, and both pesticide and personal-care product-use annotations.

**Table 7.** High-confidence DNT-IVB active chemicals with positive ToxRefDB evidence and exposure-relevant CPDat product-use context. CPDat product-use context indicates product/use association and should not be interpreted as measured exposure.

| Chemical | DNT endpoints | Min. $AC_{50}$ (μM) | Positive in vivo evidence | Product contexts | Product categories |
| --- | --- | --- | --- | --- | --- |
| Abamectin | 33 | 0.000647 | 16 | 1 | Pesticides |
| Maneb | 9 | 0.001000 | 39 | 1 | Pesticides |
| Sodium fluoride | 1 | 0.002743 | 17 | 12 | Personal care; Cleaning; Food and drug |
| Malathion | 6 | 0.002743 | 55 | 1 | Pesticides |
| Vinclozolin | 2 | 0.002786 | 73 | 1 | Pesticides |
| Sodium chlorite | 5 | 0.003000 | 7 | 4 | Personal care; Cleaning and safety |
| L-Ascorbic acid | 3 | 0.003000 | 11 | 93 | Personal care |
| Diazinon | 5 | 0.003000 | 58 | 4 | Personal care; Pesticides |
| Tris(methylphenyl) phosphate | 2 | 0.003000 | 4 | 2 | Personal care; Cleaning products |
| 2-Methoxyethanol | 1 | 0.003000 | 6 | 1 | Cleaning products |

**Figure 3.**
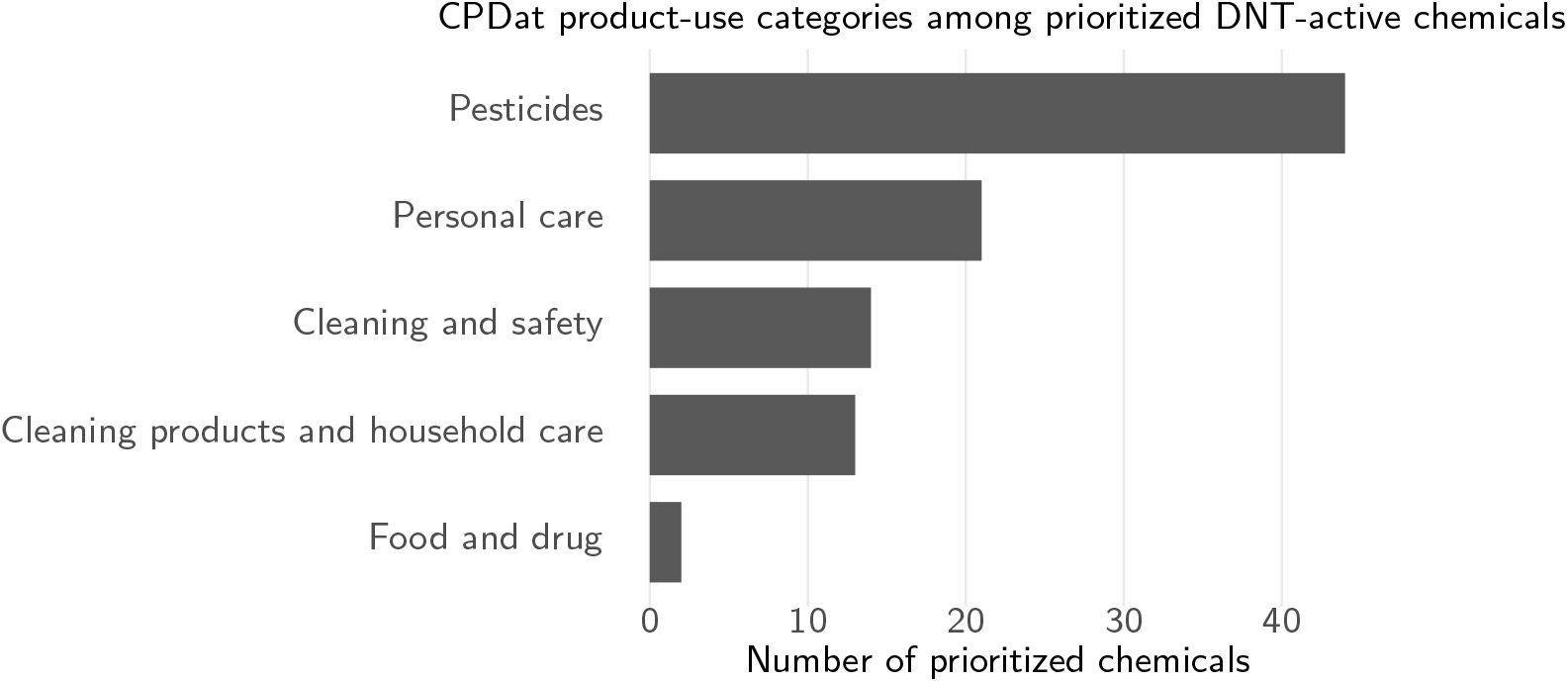
Exposure-relevant CPDat product-use categories among chemicals combining high-confidence DNT-IVB activity and positive ToxRefDB apical evidence. Bars indicate the number of distinct chemicals with at least one product-use summary in the respective CPDat category. CPDat context indicates product/use association and should not be interpreted as measured exposure.

**Figure 4.**
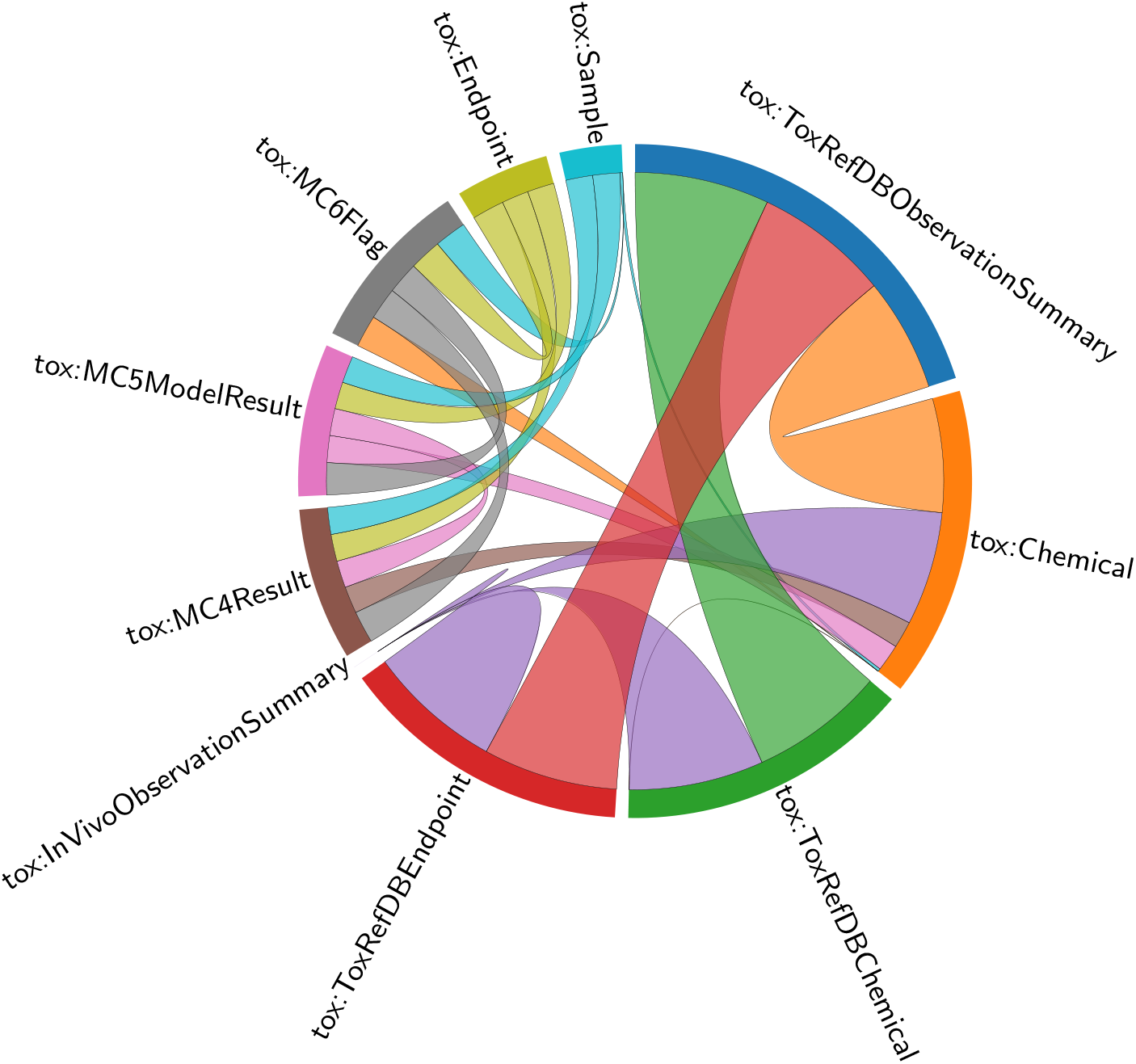
Chord diagram illustrating semantic dependencies and class relationships in the DNT-scoped ToxCast Evidence Graph with CPDat and ToxRefDB integration. Band widths are proportional to the volume of semantic links between entities.

This case study illustrates the added value of combining NAM bioactivity, in vivo apical evidence, and product-use information in a reproducible natural-language-to-SPARQL workflow. The result is not a risk assessment and does not quantify exposure magnitude. Instead, it provides a transparent prioritization view: chemicals can be ranked and interpreted according to potency, breadth of DNT-IVB activity, positive in vivo evidence, and the product-use domains in which they appear. This demonstrates how ToxCastLite can support evidence triangulation across mechanistic screening data, curated animal-study evidence, and exposure-relevant product context, thereby moving NAM-derived signals from isolated assay outputs toward integrated toxicological decision support.

## Limitations

Several limitations should be considered when interpreting ToxCastLite and the case studies presented here. First, the integrated graph is an evidence-access and prioritization system, not an automated hazard or risk-assessment engine. The queries identify chemicals for which different evidence layers co-occur, but they do not establish causality between in vitro activity, in vivo apical outcomes, and product-use context. Final interpretation remains dependent on expert review of assay biology, concentration–response behavior, study design, endpoint relevance, and data quality.

Second, the case studies should not be interpreted as formal validation of DNT-IVB assays against in vivo developmental neurotoxicity outcomes. Concordance in Case Study 1 was defined operationally as the co-occurrence of high-confidence in vitro DNT-IVB activity with positive ToxRefDB apical evidence, represented by critical-effect summaries or LOAEL point-of-departure records. This does not imply a one-to-one correspondence between individual in vitro endpoints and specific apical in vivo effects. Conversely, the absence of DNT-specific positive evidence in Case Study 2 should not be interpreted as evidence that a chemical is negative for DNT in vivo. It may reflect lack of DNT testing, incomplete endpoint coverage, study-type mismatch, curation boundaries, or true discordance between in vitro and in vivo evidence layers.

Third, the thresholds used for prioritization are pragmatic rather than definitive toxicological cutoffs. We used positive MC5 hit calls, *AC*_50_ *<* 10 *µ*M, DNT-IVB assay-list membership, and absence of MC6 quality flags to define high-confidence in vitro DNT activity. These criteria are useful for demonstrating the graph workflow, but alternative thresholds, endpoint filters, quality-flag policies, or potency metrics may be appropriate for other regulatory or scientific questions. Similarly, defining positive in vivo evidence through critical-effect summaries and LOAEL records is conservative but does not capture every possible interpretation of in vivo toxicity.

Fourth, CPDat product-use context should not be interpreted as measured exposure, exposure magnitude, or risk. CPDat records provide product-use, functional-use, list-presence, and source-document context. They can help prioritize chemicals whose bioactivity and in vivo evidence overlap with consumer, household, pesticide, or other use domains, but they do not quantify dose, frequency of use, route of exposure, population exposure, or real-world risk. Additional exposure modeling or measurement data would be required for risk characterization.

Fifth, the current RDF projection intentionally summarizes dense source data rather than materializing every source row as RDF. Detailed concentration–response records, CPDat product-composition rows, and granular ToxRefDB observation or dose-treatment group records remain in SQLite and are exposed through semantic handles, summary entities, and retrieval identifiers. This design improves portability and GraphDB queryability, but detailed review still requires drill-down into the underlying SQLite stores. Some analyses may therefore require combined SPARQL and SQL workflows.

Sixth, the graph reflects the scope, coverage, and curation status of the underlying source databases. ToxCast/invitrodb assay coverage is not uniform across chemicals and endpoints; ToxRefDB is enriched for guideline and guideline-like animal studies and has its own study-type and chemical-coverage biases; and CPDat coverage depends on available source documents and curated product-use vocabularies. Consequently, absence of evidence in the graph should be interpreted cautiously and should not be treated as evidence of absence.

Finally, the natural-language interface is intended to support query generation, not to serve as an independent source of toxicological knowledge. The local LLM acts as a schema-grounded translator from user questions to SPARQL. Generated queries should be inspected, versioned, and preserved together with their outputs when used in regulatory or scientific review. In the case studies, the generated SPARQL queries were reported in the appendix to make the workflow reproducible and to reduce ambiguity about how each natural-language question was operationalized.

## Conclusion

ToxCastLite provides a portable semantic evidence layer for integrated toxicology data mining. By combining local SQLite evidence stores with compact RDF projections, the system separates dense source-level storage from semantic discovery: SQLite preserves concentration–response records, model outputs, CPDat product-use records, and ToxRefDB in vivo study evidence, while GraphDB exposes chemicals, endpoints, assay scopes, model results, quality flags, product-use context, point-of-departure records, effect summaries, and observation summaries as queryable semantic objects.

This hybrid design allows regulatory and scientific users to ask evidence-oriented questions without directly navigating the underlying MySQL, PostgreSQL, SQLite, or RDF schemas. At the same time, the system preserves traceability by linking semantic query results back to the underlying SQLite records, including concentration–response curves, model outputs, ToxRefDB study and POD evidence, and CPDat source context for expert review. ToxCastLite therefore supports local, auditable, and reproducible exploration of NAM bioactivity, curated in vivo apical evidence, and exposure-relevant product-use context while keeping final interpretation under expert control.

The DNT case studies illustrate how the integrated graph can identify concordance between high-confidence in vitro DNT activity and positive in vivo apical evidence, highlight chemicals where in vitro DNT activity extends beyond available DNT-specific in vivo evidence, and prioritize chemicals where NAM signals, ToxRefDB evidence, and CPDat product-use context intersect. These examples show how semantic evidence integration can move NAM-derived signals from isolated screening outputs toward transparent, context-rich evidence navigation.

The approach is intended as a practical evidence-access framework rather than an automated decision system. It can support prioritization, review, evidence-gap analysis, and hypothesis generation in regulatory toxicology workflows where data sovereignty, transparency, and drill-down access to source-level evidence are essential.

## Data and responsibility statement

The analyses presented in this work are based on publicly available source data from ToxCast/invitrodb, ToxRefDB, and CPDat. ToxCastLite generates derived SQLite evidence stores and RDF projections to support semantic access, evidence linking, prioritization, and drill-down, but it does not modify the underlying scientific content of the source databases. The authors make no warranty regarding the completeness, accuracy, regulatory sufficiency, or fitness for a specific decision-making purpose of the underlying source data or of the derived evidence summaries. Results should be interpreted in light of the coverage, curation status, and limitations of the original data resources, and final toxicological or regulatory conclusions remain the responsibility of qualified experts.

## Author contributions statement

A.D. conceived the project, designed the architecture, implemented the SQLite extraction pipeline, designed and implemented the RDF projection workflow, developed the Streamlit prototype, performed database builds and validation queries, interpreted the results, and drafted the manuscript. O.N. contributed to the local deployment of Gemma 4 and Qwen 3.5 within the institutional infrastructure and, together with K.H., tested and validated example questions and query workflows. A.M. provided scientific bioinformatics advice. K.H., E.F. and K.K. provided scientific toxicological advice.

## Appendix

### Chord diagram

### Generated SPARQL queries

The following SPARQL queries were generated from the natural-language questions entered in the Streamlit interface by the local LLM and executed against the integrated ToxCastLite graph, including ToxCast bioactivity, ToxRefDB in vivo evidence, and CPDat product-use context where applicable.

#### Case Study 1: Concordance between in vitro DNT activity and in vivo apical evidence

~~~
PREFIX tox: <https://w3id.org/toxcast-evidence/vocab/>
PREFIX rdfs: <http://www.w3.org/2000/01/rdf-schema#>
PREFIX xsd: <http://www.w3.org/2001/XMLSchema#>
SELECT
 ?chemical_label
 (COUNT(DISTINCT ?endpoint) AS ?n_active_dnt_endpoints)
 (MIN(xsd:decimal(?ac50)) AS ?min_ac50_uM)
 (COUNT(DISTINCT ?positive_in_vivo_evidence) AS ?n_positive_in_vivo_evidence)
 (COUNT(DISTINCT ?critical_effect_summary) AS ?n_critical_effect_summaries)
 (COUNT(DISTINCT ?loael_pod) AS ?n_loael_pods)
 (GROUP_CONCAT(DISTINCT ?study_type; separator=“ | ”) AS ?toxrefdb_study_types)
 (GROUP_CONCAT(DISTINCT ?evidence_type; separator=“ | ”) AS ?positive_evidence_types)
 (GROUP_CONCAT(DISTINCT ?endpoint_category; separator=“ | ”) AS ?in_vivo_endpoint_categories)
 (GROUP_CONCAT(DISTINCT ?toxval_effect_list; separator=“ | ”) AS ?toxval_effects)
WHERE {
 # High-confidence in vitro DNT-IVB activity
 ?mc5 a tox:MC5ModelResult ;
 tox:hitCall ?hit_call ;
 tox:ac50 ?ac50 ;
 tox:forChemical ?chemical ;
 tox:forEndpoint ?endpoint .
 FILTER(xsd:decimal(?hit_call) > 0)
 FILTER(xsd:decimal(?ac50) < 10)
 # Exclude MC5 model results with MC6 quality flags
 FILTER NOT EXISTS {?mc5 tox:hasFlag ?flag .}
 FILTER NOT EXISTS {?mc5 tox:hasFlagValue ?flag_value .}
 ?endpoint tox:belongsToAssayList ?assay_list .
 ?assay_list rdfs:label ?assay_list_label .
 FILTER(LCASE(STR(?assay_list_label)) = “dnt-ivb”)
 ?chemical rdfs:label ?chemical_label .
 # Positive in vivo apical evidence from ToxRefDB
 {
 ?chemical tox:hasToxRefDBEffectSummary ?critical_effect_summary .
 ?critical_effect_summary
 a tox:ToxRefDBEffectSummary ;
 tox:criticalEffectCount ?n_critical ;
 tox:studyType ?study_type .
 FILTER(xsd:integer(?n_critical) > 0)
 FILTER(?study_type IN (“DNT”, “DEV”, “MGR”, “CHR”, “SUB”))
 OPTIONAL {
 ?critical_effect_summary tox:endpointCategory ?endpoint_category .
 }
 BIND(?critical_effect_summary AS ?positive_in_vivo_evidence)
 BIND(“critical effect summary” AS ?evidence_type)
}
 UNION
 {
 ?chemical tox:hasToxRefDBPointOfDeparture ?loael_pod .
 ?loael_pod
 a tox:ToxRefDBPointOfDeparture ;
 tox:calcPodType ?pod_type ;
 tox:studyType ?study_type .
 FILTER(STR(?pod_type) = “LOAEL”)
 FILTER(?study_type IN (“DNT”, “DEV”, “MGR”, “CHR”, “SUB”))
 OPTIONAL {
 ?loael_pod tox:toxvalEffectList ?toxval_effect_list .
 }
 BIND(?loael_pod AS ?positive_in_vivo_evidence)
 BIND(“LOAEL POD” AS ?evidence_type)
}
}
GROUP BY ?chemical_label
ORDER BY ?min_ac50_uM DESC(?n_active_dnt_endpoints) DESC(?n_positive_in_vivo_evidence)
~~~

#### Case Study 2: In vitro DNT activity beyond available in vivo DNT evidence

~~~
PREFIX tox: <https://w3id.org/toxcast-evidence/vocab/>
PREFIX rdfs: <http://www.w3.org/2000/01/rdf-schema#>
PREFIX xsd: <http://www.w3.org/2001/XMLSchema#>
SELECT
 ?chemical_label
 (COUNT(DISTINCT ?endpoint) AS ?n_active_dnt_endpoints)
 (MIN(xsd:decimal(?ac50)) AS ?min_ac50_uM)
 (COUNT(DISTINCT ?study) AS ?n_toxrefdb_studies)
 (COUNT(DISTINCT ?positive_non_dnt_evidence) AS ?n_positive_non_dnt_evidence)
 (GROUP_CONCAT(DISTINCT ?study_type; separator=“ | ”) AS ?available_toxrefdb_study_types)
 (GROUP_CONCAT(DISTINCT ?positive_non_dnt_study_type; separator=“ | ”) AS ?positive_non_dnt_study_types)
 (GROUP_CONCAT(DISTINCT ?endpoint_category; separator=“ | ”) AS ?positive_non_dnt_endpoint_categories)
WHERE {
 # High-confidence in vitro DNT-IVB activity
 ?mc5 a tox:MC5ModelResult ;
 tox:hitCall ?hit_call ;
 tox:ac50 ?ac50 ;
 tox:forChemical ?chemical ;
 tox:forEndpoint ?endpoint .
 FILTER(xsd:decimal(?hit_call) > 0)
 FILTER(xsd:decimal(?ac50) < 10)
 # Exclude MC5 model results with MC6 quality flags
 FILTER NOT EXISTS {?mc5 tox:hasFlag ?flag .}
 FILTER NOT EXISTS {?mc5 tox:hasFlagValue ?flag_value .}
 ?endpoint tox:belongsToAssayList ?assay_list .
 ?assay_list rdfs:label ?assay_list_label .
 FILTER(LCASE(STR(?assay_list_label)) = “dnt-ivb”)
 ?chemical rdfs:label ?chemical_label .
 # Require available processed ToxRefDB study evidence
 ?chemical tox:hasToxRefDBStudy ?study .
 ?study tox:studyType ?study_type .
 FILTER(?study_type IN (“DNT”, “DEV”, “MGR”, “CHR”, “SUB”))
 # Exclude chemicals with DNT-specific positive in vivo evidence.
 # Positive DNT evidence is defined as either:
 # - DNT critical-effect summary with criticalEffectCount > 0
 # - DNT LOAEL POD
 FILTER NOT EXISTS {
 {
 ?chemical tox:hasToxRefDBEffectSummary ?dnt_effect_summary .
 ?dnt_effect_summary
 a tox:ToxRefDBEffectSummary ;
 tox:studyType “DNT” ;
 tox:criticalEffectCount ?n_dnt_critical .
 FILTER(xsd:integer(?n_dnt_critical) > 0)
 }
 UNION
 {
 ?chemical tox:hasToxRefDBPointOfDeparture ?dnt_loael_pod .
 ?dnt_loael_pod
 a tox:ToxRefDBPointOfDeparture ;
 tox:studyType “DNT” ;
 tox:calcPodType ?dnt_pod_type .
 FILTER(STR(?dnt_pod_type) = “LOAEL”)
 }
}
# Optional: capture positive non-DNT in vivo evidence.
# This shows whether the chemical has positive apical evidence elsewhere,
# even if DNT-specific positive evidence is absent.
OPTIONAL {
 {
 ?chemical tox:hasToxRefDBEffectSummary ?positive_non_dnt_evidence .
 ?positive_non_dnt_evidence
 a tox:ToxRefDBEffectSummary ;
 tox:studyType ?positive_non_dnt_study_type ;
 tox:criticalEffectCount ?n_critical .
 FILTER(xsd:integer(?n_critical) > 0)
 FILTER(?positive_non_dnt_study_type IN (“DEV”, “MGR”, “CHR”, “SUB”))
 OPTIONAL {
 ?positive_non_dnt_evidence tox:endpointCategory ?endpoint_category .
 }
}
UNION
{
 ?chemical tox:hasToxRefDBPointOfDeparture ?positive_non_dnt_evidence .
 ?positive_non_dnt_evidence
 a tox:ToxRefDBPointOfDeparture ;
 tox:studyType ?positive_non_dnt_study_type ;
 tox:calcPodType ?pod_type .
 FILTER(STR(?pod_type) = “LOAEL”)
 FILTER(?positive_non_dnt_study_type IN (“DEV”, “MGR”, “CHR”, “SUB”))
 }
}
}
GROUP BY ?chemical_label
ORDER BY ?min_ac50_uM DESC(?n_active_dnt_endpoints) DESC(?n_positive_non_dnt_evidence)
~~~

#### Case Study 3: Exposure-relevant prioritization across NAM, in vivo, and product-use evidence

~~~
PREFIX tox: <https://w3id.org/toxcast-evidence/vocab/>
PREFIX rdfs: <http://www.w3.org/2000/01/rdf-schema#>
PREFIX xsd: <http://www.w3.org/2001/XMLSchema#>
SELECT
 ?chemical_label
 ?n_active_dnt_endpoints
 ?min_ac50_uM
 ?n_positive_in_vivo_evidence
 ?n_product_contexts
 ?total_distinct_products
 ?toxrefdb_study_types
 ?positive_evidence_types
 ?in_vivo_endpoint_categories
 ?product_categories
 ?product_contexts
WHERE {
 # ----------------------------------------------------------
 # High-confidence in vitro DNT-IVB activity
 # ----------------------------------------------------------
{
 SELECT
 ?chemical
 ?chemical_label
 (COUNT(DISTINCT ?endpoint) AS ?n_active_dnt_endpoints)
 (MIN(xsd:decimal(?ac50)) AS ?min_ac50_uM)
 WHERE {
 ?mc5 a tox:MC5ModelResult ;
 tox:hitCall ?hit_call ;
 tox:ac50 ?ac50 ;
 tox:forChemical ?chemical ;
 tox:forEndpoint ?endpoint .
 FILTER(xsd:decimal(?hit_call) > 0)
 FILTER(xsd:decimal(?ac50) < 10)
 # Exclude MC5 model results with MC6 quality flags
 FILTER NOT EXISTS {?mc5 tox:hasFlag ?flag .}
 FILTER NOT EXISTS {?mc5 tox:hasFlagValue ?flag_value .}
 ?endpoint tox:belongsToAssayList ?assay_list .
 ?assay_list rdfs:label ?assay_list_label .
 FILTER(LCASE(STR(?assay_list_label)) = “dnt-ivb”)
 ?chemical rdfs:label ?chemical_label .
 }
 GROUP BY ?chemical ?chemical_label
}
# ----------------------------------------------------------
# Positive in vivo apical evidence from ToxRefDB
# Positive = critical-effect summary OR LOAEL POD
# ----------------------------------------------------------
{
 SELECT
 ?chemical
 (COUNT(DISTINCT ?positive_in_vivo_evidence) AS ?n_positive_in_vivo_evidence)
 (GROUP_CONCAT(DISTINCT ?study_type; separator=“ | ”) AS ?toxrefdb_study_types)
 (GROUP_CONCAT(DISTINCT ?evidence_type; separator=“ | ”) AS ?positive_evidence_types)
 (GROUP_CONCAT(DISTINCT ?endpoint_category; separator=“ | ”) AS ?in_vivo_endpoint_categories)
 WHERE {
 {
 ?chemical tox:hasToxRefDBEffectSummary ?positive_in_vivo_evidence .
 ?positive_in_vivo_evidence
 a tox:ToxRefDBEffectSummary ;
 tox:criticalEffectCount ?n_critical ;
 tox:studyType ?study_type .
 FILTER(xsd:integer(?n_critical) > 0)
 FILTER(?study_type IN (“DNT”, “DEV”, “MGR”, “CHR”, “SUB”))
 OPTIONAL {
 ?positive_in_vivo_evidence tox:endpointCategory ?endpoint_category .
 }
 BIND(“critical effect summary” AS ?evidence_type)
 }
 UNION
 {
 ?chemical tox:hasToxRefDBPointOfDeparture ?positive_in_vivo_evidence .
 
 ?positive_in_vivo_evidence
 a tox:ToxRefDBPointOfDeparture ;
 tox:calcPodType ?pod_type ;
 tox:studyType ?study_type .
 FILTER(STR(?pod_type) = “LOAEL”)
 FILTER(?study_type IN (“DNT”, “DEV”, “MGR”, “CHR”, “SUB”))
 OPTIONAL {
 ?positive_in_vivo_evidence tox:toxvalEffectList ?endpoint_category .
 }
 
 BIND(“LOAEL POD” AS ?evidence_type)
 }
}
GROUP BY ?chemical
}
# ----------------------------------------------------------
# CPDat product-use context
# Exposure-relevant contexts:
# children/toys, personal care, household/cleaning, pesticides
# ----------------------------------------------------------
{
 SELECT
 ?chemical
 (COUNT(DISTINCT ?pus) AS ?n_product_contexts)
 (SUM(xsd:integer(?n_products_raw)) AS ?total_distinct_products)
 (GROUP_CONCAT(DISTINCT ?puc_general_category; separator=“ | ”) AS ?product_categories)
 (GROUP_CONCAT(DISTINCT ?product_context; separator=“ | ”) AS ?product_contexts)
 WHERE {
 ?chemical tox:hasProductUseSummary ?pus .
 ?pus tox:pucGeneralCategory ?puc_general_category ;
 tox:pucProductFamily ?puc_product_family .
 OPTIONAL {?pus tox:pucProductType ?puc_product_type .}
 OPTIONAL {?pus tox:distinctProductCount ?n_products_raw .}
 FILTER(STR(?puc_general_category) != “(missing)”)
 FILTER(STR(?puc_product_family) != “(missing)”)
 FILTER(
 CONTAINS(LCASE(STR(?puc_general_category)), “child”) ||
 CONTAINS(LCASE(STR(?puc_product_family)), “child”) ||
 CONTAINS(LCASE(STR(?puc_product_type)), “child”) ||
 CONTAINS(LCASE(STR(?puc_general_category)), “toy”) ||
 CONTAINS(LCASE(STR(?puc_product_family)), “toy”) ||
 CONTAINS(LCASE(STR(?puc_product_type)), “toy”) ||
 CONTAINS(LCASE(STR(?puc_general_category)), “personal care”) ||
 CONTAINS(LCASE(STR(?puc_product_family)), “personal care”) ||
 CONTAINS(LCASE(STR(?puc_general_category)), “cleaning”) ||
 CONTAINS(LCASE(STR(?puc_product_family)), “cleaning”) ||
 CONTAINS(LCASE(STR(?puc_general_category)), “household”) ||
 CONTAINS(LCASE(STR(?puc_product_family)), “household”) ||
 CONTAINS(LCASE(STR(?puc_general_category)), “pesticide”) ||
 CONTAINS(LCASE(STR(?puc_product_family)), “pesticide”)
)
 BIND(
 IF(
 BOUND(?puc_product_type) && STR(?puc_product_type) != “(missing)”,
 CONCAT(STR(?puc_product_family), “ / ”, STR(?puc_product_type)),
 STR(?puc_product_family)
)
 AS ?product_context
)
 }
 GROUP BY ?chemical
 }
}
ORDER BY
 ?min_ac50_uM
 DESC(?n_active_dnt_endpoints)
 DESC(?n_positive_in_vivo_evidence)
 DESC(?n_product_contexts)
~~~

## References

[1] Ann M Richard, Richard S Judson, Keith A Houck, Christopher M Grulke, Patra Volarath, Inthirany Thillainadarajah, Chihae Yang, James Rathman, Matthew T Martin, John F Wambaugh, et al. Toxcast chemical landscape: paving the road to 21st century toxicology. Chemical research in toxicology, 29(8):1225–1251, 2016.

[2] David J Dix, Keith A Houck, Matthew T Martin, Ann M Richard, R Woodrow Setzer, and Robert J Kavlock. The toxcast program for prioritizing toxicity testing of environmental chemicals. Toxicological sciences, 95(1):5–12, 2007.

[3] EPA’s Center for Computational Toxicology and Exposure. ToxCast Database: invitrodb version 4.3. 5 2018.

[4] Renzo Angles and Claudio Gutierrez. Survey of graph database models. ACM computing surveys (CSUR), 40(1):1–39, 2008.

[5] EPA’s Center for Computational Toxicology and Exposure. ToxRefDB v3.0, 3 2018.

[6] Madison Feshuk, Lori Kolaczkowski, Sean Watford, and Katie Paul Friedman. Toxrefdb v2.1: update to curated in vivo study data in the toxicity reference database. Frontiers in Toxicology, 5:1260305, 2023.

[7] Thomas Sheffield, Jason Brown, Sarah Davidson, Katie Paul Friedman, and Richard Judson. tcplfit2: an r-language general purpose concentration– response modeling package. Bioinformatics, 38(4):1157–1158, 2022.

[8] Sakshi Handa, Kristin K Isaacs, Jonathan T Wall, Allison Larger, Scott Burns, Lauren E Koval, Kenta Baron-Furuyama, Colleen M Elonen, David Lyons, Kathie L Dionisio, et al. The chemical and products database v4. 0, an updated resource supporting chemical exposure evaluations. Scientific Data, 12(1):950, 2025.

